# High-affinity low-capacity and low-affinity high-capacity N-acetyl-2-aminofluorene (AAF) macromolecular binding sites are revealed during the growth cycle of adult rat hepatocytes in primary culture

**DOI:** 10.1101/209130

**Authors:** K.S. Koch, T. Moran, W.T. Shier, H.L. Leffert

## Abstract

Long-term cultures of primary adult rat hepatocytes were used to study the effects of N-acetyl-2-aminofluorene (AAF) on hepatocyte proliferation during the growth cycle; on the initiation of hepatocyte DNA synthesis in quiescent cultures; and, on hepatocyte DNA replication following the initiation of DNA synthesis. Scatchard analyses were used to identify the pharmacologic properties of radiolabeled AAF metabolite binding to hepatocyte macromolecules. Two classes of growth cycle-dependent AAF metabolite binding sites – a high-affinity low-capacity site (designated Site I) and a low-affinity high-capacity site (designated Site II) – associated with two spatially distinct classes of macromolecular targets, were revealed. Based upon radiolabeled AAF metabolite binding to purified hepatocyte genomic DNA or to DNA, RNA, proteins and lipids from isolated nuclei, Site I_DAY 4_ targets (K_D[APPARENT]_ ≈ 2-4 x 10^−6^ M and B_MAX[APPARENT]_ ≈ 6 pmols/10^6^ cells/24 h) were consistent with genomic DNA; and with AAF metabolized by a nuclear cytochrome P450. Based upon radiolabeled AAF binding to total cellular lysates, Site II_DAY 4_ targets (K_D[APPARENT]_ ≈ 1.5 x 10^−3^ M and B_MAX[APPARENT]_ ≈ 350 pmols/10^6^ cells/24 h) were consistent with cytoplasmic proteins; and with AAF metabolized by cytoplasmic cytochrome P450s. DNA synthesis was not inhibited by concentrations of AAF that saturated DNA binding in the neighborhood of the Site I K_D_. Instead, hepatocyte DNA synthesis inhibition required higher concentrations of AAF approaching the Site II K_D_. These observations raise the possibility that carcinogenic DNA adducts derived from AAF metabolites form below concentrations of AAF that inhibit replicative and repair DNA synthesis.

## INTRODUCTION

N-acetyl-2-aminofluorene (AAF) is a well-established hepatoprocarcinogen; its chemical activation pathways and toxicity have been studied in detail (Wilson *et al.*, 1941; Miller *et al.*, 1960; Weisburger and Weisburger, 1973). Less is known about the biological effects, cellular attributes and pharmacology of AAF binding to macromolecules inside living hepatocytes. To address these problems, a series of experiments have been performed using a well-characterized, proliferation-competent, long-term primary adult rat hepatocyte culture system (Leffert and Koch, 1977; Leffert *et al.*, 1977, 1978a,b; Koch and Leffert, 1979, 1980; Leffert *et al.*, 1983; Koch *et al.*, 1982; submitted, 2017)^1^.

The investigations described here comprise the second report of two, the first of which described investigations focused upon AAF metabolism (Koch *et al.*, submitted, 2017). In the first report, Lineweaver-Burk analyses of AAF metabolites produced by hepatocytes revealed two principal classes of hepatocyte AAF metabolism: System I (high-affinity and low-velocity), K_m[APPARENT]_ = 1.64 x 10^−7^ M and V_MAX[APPARENT]_ « 0.1 nmols/10^6^ cells/24 h; and, System II (low-affinity and high-velocity), K_m[APPARENT]_ = 3.25 x 10^−5^ M and V_MAX[APPARENT]_ « 1000 nmols/10^6^ cells/24 h. These findings and autoradiographic observations suggested selective roles and intracellular locations for System I- and System II-mediated AAF metabolite formation inside hepatocytes.

In this report, experiments were designed to elucidate and quantify the intracellular properties of the binding of AAF metabolites with respect to hepatocyte proliferation, the initiation of hepatocyte DNA synthesis, and hepatocyte DNA replication. The results reveal several unrecognized aspects of AAF metabolite binding in primary hepatocyte cultures. As with the patterns of AAF metabolism, hepatocytes exhibit two classes of macromolecular AAF metabolite binding sites: Site I (high-affinity and low-capacity); and, Site II (low-affinity and high-capacity). The binding of metabolites of AAF to these sites differentially affects the inhibition of hepatocyte growth and macromolecular synthesis. Scatchard analyses provide further evidence of the growth cycle dependence of the levels of Site I and Site II constants. Taken together, the results have important implications for mechanisms of chemical hepatocarcinogenesis, even if non-parenchymal or liver stem cells are adduct-forming bystanders and/or carcinogen targets (Koch and Leffert, 2015).

## MATERIALS AND METHODS

### Sources of radiolabeled compounds

N-acetyl-9-[^14^C]-2-aminofluorene ([^14^C]-AAF) was purchased from NEN (New England Nuclear, Boston, MA [Leffert *et al.*, 1977; Koch *et al.*, submitted, 2017]). The specific activity of [^14^C]-AAF was 100 dpm pmol^−1^. [Ring G-^3^H]-N-acetyl-2-aminofluorene ([^3^H]-AAF) was synthesized from 2-aminofluorene (AF) and purified as described elsewhere (Koch *et al.*, submitted, 2017). The specific activity of [^3^H]-AAF was 4000 dpm pmol^−1^.

### Primary hepatocyte culture

Hepatocytes were isolated from adult male Fischer 344 or lipotrope-deficient (Rogers, 1975) Sprague Dawley rats (180–200 g) by perfusion through the portal vein with a collagenase cocktail, and plated as described previously (Leffert *et al.*, 1979). All measurements were made using 3 dishes per point per plating with errors ± 10%. In experiments comprised of multiple animals, the hepatocytes from each animal were plated separately, and each point of the results consists of multiple sets of 3 dish averages, unless otherwise stated.

### Primary hepatocyte growth initiation assays

Growth initiation assays (DNA synthesis, S-phase entry and mitosis) were performed with 12 day-old quiescent (G_0_) cultures (≈ 8 x 10^5^ cells/dish) as described (Koch and Leffert, 1979). Control cultures were fluid-changed into fresh media containing 50 ng each of insulin, glucagon and epidermal growth factor (EGF)/mL, supplemented with AAF vehicle (1% v/v EtOH). Experimental cultures were shifted into the same medium to which varying concentrations of AAF were added at zero time or 12 h later. DNA synthesis was measured by the uptake of 1.25 μCi [^3^H]-TdR/mL (3 μM TdR) into trichloroacetic acid (TCA)-insoluble cpm/10^6^ cells, at various times after the medium changes (Koch and Leffert, 1979). Three μM TdR saturates the Km of the TdR transport system which obviates artifacts of growth factor induced changes in dTTP pool sizes (Paul *et al.*, 1972).

### Quantification of cellular levels of intracellular free AAF ([AAF]_i_) and covalently bound AAF metabolites

At various times post-plating, [^3^H]-AAF or [^14^C]-AAF were added to the cultures for different intervals of time, as indicated in Results and Legends. The monolayers were washed 3x with 2 mL of ice-cold Ca^2+^- and Mg^2+^-supplemented Tris-HCl buffer, pH 7.4, and extracted with 2 mL of 5% v/v TCA overnight at 4°C in darkness. The soluble acid extracts were subjected to liquid scintillation counting (Leffert *et al.*,1979), the results of which were used to quantify the cellular concentrations of free [AAF]_i_. Intracellular spaces were found to be 2 pL H2O/cell (Koch and Leffert, 1979); it was assumed that [AAF]_i_ distributed into this space. The extracted monolayers were washed 3x with Tris-HCl buffer, pH 7.4, solubilized (Kruijer *et al.*, 1986), heated at 100°C for 2 min, and brought to room temperature. Covalently bound radiolabeled fluorene residues in these extracts were precipitated with ice-cold 10% TCA onto Whatman GF/C™ filters (Sigma Aldrich). The filters were washed with ice-cold EtOH, air dried, and subjected to scintillation counting to errors of ± 1%. Counting efficiencies of [^14^C] and [^3^H] standards were 90-95% and 50-55%, respectively.

### DNA isolation and quantification, and isolation of nuclei from cultured hepatocytes

Purified RNA-free deproteinized genomic DNA was isolated from the nuclei (Blobel and Potter, 1966) of cultured primary rat hepatocytes, using a modified Marmur method (Paul and Gilmour, 1968), and quantified by the Burton procedure (Waterborg and Matthews, 1985).

### Autoradiography

Following medium changes, 12 day-old cultures were labeled 12-24 h later with 1.25 μCi [^3^H]-TdR/mL (3 μM TdR). At 24 h, the cultures were washed 6x with Tris-HCl buffer, pH 7.4; fixed with neutral buffered formalin; exposed to β-particle sensitive Eastman Kodak™ AR-10 stripping film, developed and stained with crystal violet, as described elsewhere (Koch and Leffert, 1974).

### Statistical analyses

Linear regression curves and Scatchard analyses were calculated by standard procedures using Prism™ and GraphPad™ Software, Inc., La Jolla, CA.

## RESULTS

### AAF inhibits hepatocyte proliferation during the growth cycle

**Figure 1A** shows a set of growth cycle curves during an interval of 0-10 days postplating. Compared to the AAF vehicle (1 % v/v EtOH), a pulsed 24 h exposure to AAF between 1–2 days post-plating reduced both the rates of log phase proliferation, starting between 3–4 days post-plating, and the levels of stationary phase attained between 6–10 days post-plating. Growth inhibition was proportional to the initial concentrations of extracellular AAF ([AAF]_o_), which ranged between 6 x 10^−8^ M − 2 x 10^−5^ M. Detached floating cells were not observed, consistent with the absence of AAF cytotoxicity under these conditions (Howard *et al.*, 1981).

**Figure 1.**
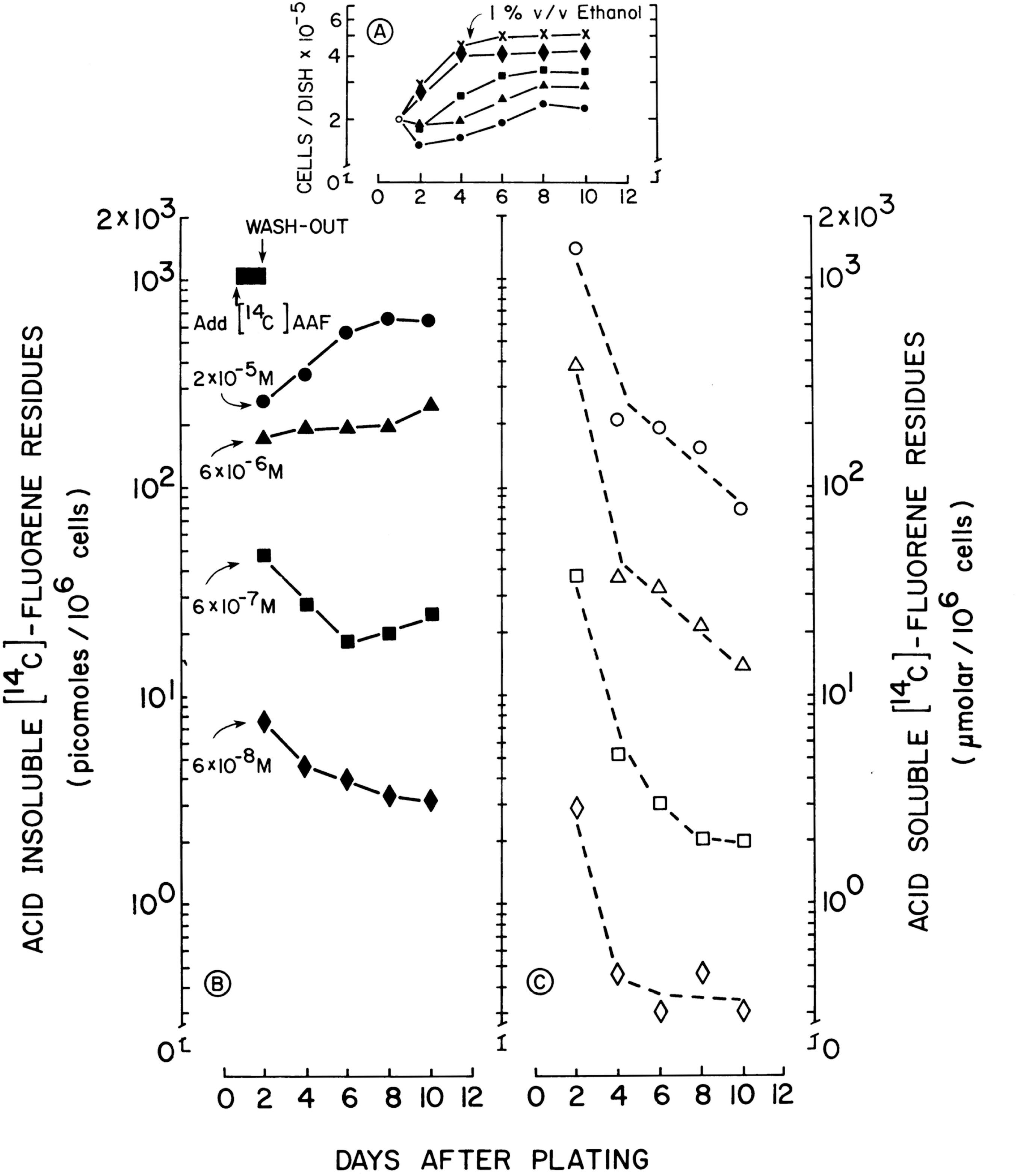
Effects of AAF on cell proliferation, binding and uptake during the growth cycle. Hepatocytes were plated from Fischer 344 rats. [^14^C]-AAF was added to the cultures (6 x 10^−8^ M, 6 x 10^−7^M, 6 x 10^−6^ M or 2 x 10^−5^ M) one day post-plating. Twenty-four h later, the dishes were washed 3x with serum-free media. Next, centrifuged 2 day-old conditioned medium (Koch and Leffert, 1974) from untreated sister cultures was added to the washed dishes, and measurements were made every 2 days through day 10 post-plating (N=3 dishes/point, with errors ± 10%). (**A**), (**B**) and (**C**), Days after plating (x-axes). (**A**) Population growth curves (top panel; vehicle, 1% v/v EtOH [**X—X**]); (**B**) Acid-insoluble [^14^C]-fluorene residues (picomoles/10^6^ cells, left y-axis); and, (**C**) Acid-soluble [^14^C]-[AAF]_i_ (μmolar/10^6^ cells, right y-axis). Symbols are all the same in all 3 panels (see annotated arrows in (**B**)).

### Changes in acid-insoluble [^14^C]-AAF binding and acid-soluble [^14^C]-[AAF]_i_ during the growth cycle

AAF exposure conditions were similar to those used in **Figure 1A**. Under these conditions, during the hepatocyte growth cycle, the levels of acid-insoluble AAF metabolite binding varied disproportionately with respect to the concentrations of pulsed [AAF]_o_ (**Figure 1B**). The shapes of the curves of covalently bound acid-insoluble [^14^C]-fluorene residues followed rising growth curve elevations at AAF concentrations of 2 x 10^−5^ M and 6 x 10^−6^ M; and, a U-shaped curve at 6 x 10^−7^ M AAF. At 6 x 10^−8^ M AAF, the lowest AAF exposure, the curve fell continuously to 38%of its initial level. In contrast, in parallel growth cycle measurements with sister cultures, when extracellular concentrations of pulsed [AAF]_o_ were reduced from 2 x 10^−5^ M to 6 x 10^−8^ M, acid-soluble [AAF]_i_ [^14^C]-fluorene residues fell proportionately between days 2-10 to 3-5% of their initial day 2 levels (**Figure 1C**). The disproportional patterns of the binding curves were probably due to the regulatory limits set by the K_m[APPARENT]_ value of AAF metabolism (1-3 x 10^−7^ M [see Koch *et al.*, submitted, 2017]). For example, AAF uptake and metabolism were not limited when concentrations of 2 x 10^−5^ M and 6 x 10^−6^ M [AAF]_o_ exceeded the Site I K_m[APPARENT]_; but, AAF uptake and metabolism became limiting when concentrations of 6 x 10^−7^ M and 6 x 10^−8^ M AAF fell below both the Site I and the Site II K_m[APPARENT]_ (≈ 2-5 x 10^−5^ M [see Koch *et al.*, submitted, 2017]). This interpretation also coincides with the finding that production of N-OH-AAF fell with 1^st^ order kinetics during the growth cycle (Koch *et al.*, submitted, 2017), which would have attenuated the pool sizes of activated electrophiles available for covalent binding.

### Effects of AAF on the initiation of DNA synthesis and DNA replication

Over a 4-log concentration range (2 x 10^−7^ M − 2 x 10^−4^ M), AAF had only a slight effect on [^3^H]-TdR uptake into an intracellular acid-soluble pool (**Figure 2A**), eliminating significant artifacts of [^3^H]-TdR transport inhibition by AAF. The initiation of DNA synthesis in 12 day-old G0 cultures was inhibited > 80% at an AAF concentration of 2 x 10^−4^ M, whereas relatively little inhibition of DNA synthesis was observed at AAF concentrations ≤ 2 x 10^−5^ M (Figure 2B). Basal DNA synthesis rates (no fresh media changes) were significantly less susceptible to inhibition by AAF (Figure 2B), as would be expected from the small fraction of cycling cells in G0 cultures (Koch and Leffert, 1979). Cellular detachment was unaffected over the 4-log AAF concentration range, excluding cytotoxicity as an underlying cause of the inhibitory changes (Howard *et al.*,1981).

**Figure 2.**
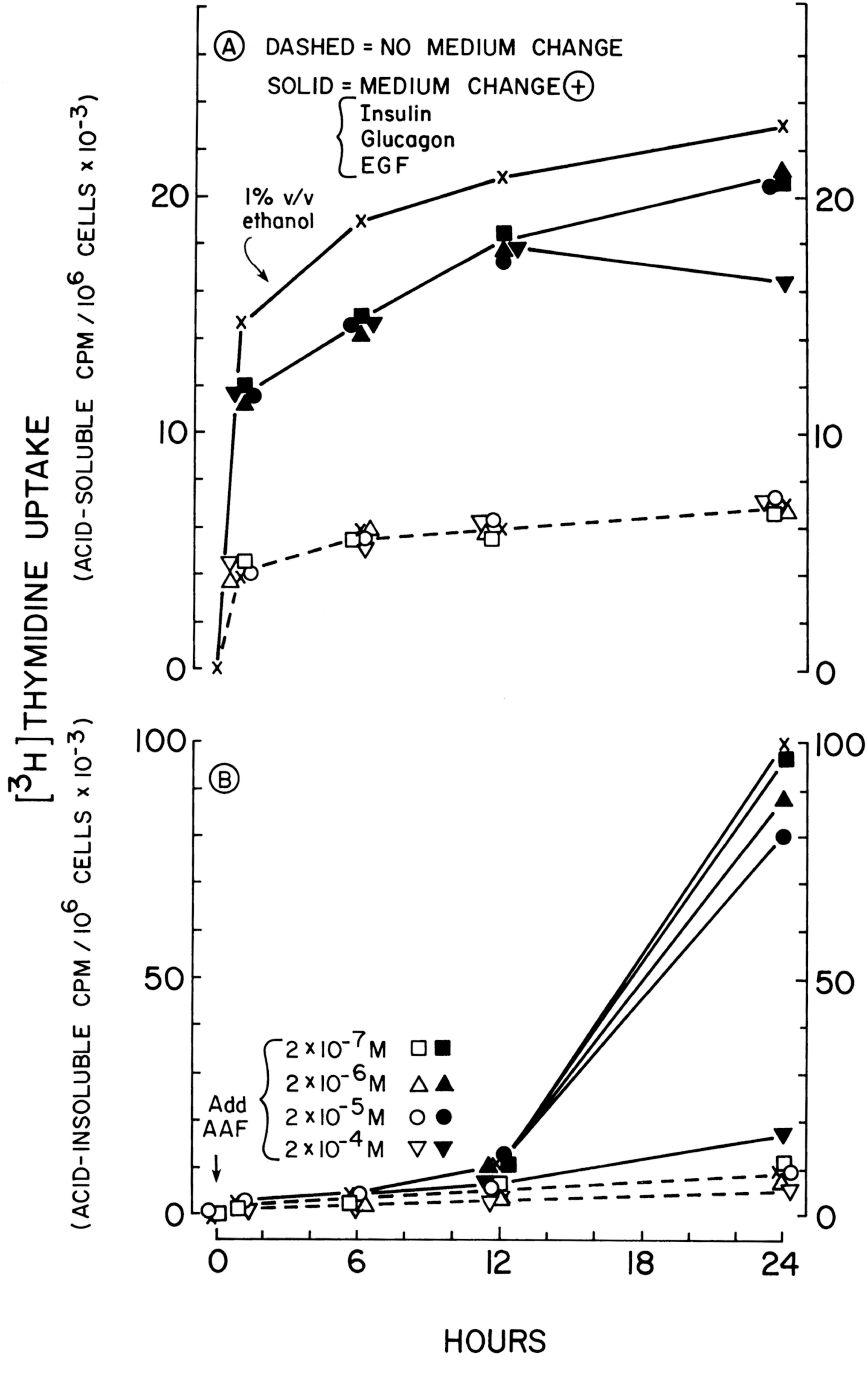
Effects of AAF on growth initiation of quiescent hepatocytes. Liver cell cultures were plated from Fischer 344 rats. Unlabeled AAF was added when indicated by the arrow (↓) over a broad concentration range (2 x 10^−7^ M, 2 x 10^−6^ M, 2 x 10^−5^ M or 2 x 10^−4^ M) to 12 day-old hepatocyte cultures (dotted lines), or to cultures fluid-changed into fresh growth-inducing media containing 50 ng each of insulin, glucagon and EGF/mL (solid lines). Before harvesting, the cultures were pulse-labeled for 4 h with 1.25 μCi [^3^H]-TdR/mL (3 μM TdR), and at the indicated times after the fluid changes (Hours [x-axes]), [^3^H]-TdR uptake was measured (y-axes). **(A)** Acid-soluble [^3^H] radioactivity (cpm/10^6^ cells x 10^−3^ (top panel); **(B)** Acid-insoluble [^3^H] radioactivity (cpm/10^6^ cells x 10^−3^ (bottom panel). The treatment symbols annotated in **(B)** are identical in **(A)**.

When 2 x 10^−4^ M AAF was added 12 h after growth initiation, DNA synthesis was still inhibited (data not shown). This suggested that this high concentration of AAF was sufficient for inhibition of mitogen stimulated DNA replication at the start of or during S-phase (Koch and Leffert, 1979; Zurlo *et al.*, 1986); but did not preclude concomitant inhibition of initiation of DNA synthesis by the 0-24 h exposure, or exclude the also possible sufficiency of a 0-12 h, high AAF exposure to inhibit initiation, and thereby subsequent progression beyond G_0_. Notably, 24 h of continuous exposure to 2 x 10^−4^ M AAF rendered the cells refractory to new mitogens in AAF-free sister culture conditioned medium over the next 24 h interval (data not shown). Taken together, these observations suggested that the initial prolonged exposure to AAF continued to block the formation or activation of one or more growth regulatory molecules, the intracellular concentrations of which were rate-limiting for DNA synthesis at cell-cycle transition points during G_0,1_→S→G_2_→M transitions (Zurlo *et al.*, 1986; Ohlson *et al.*,1998).

The inhibition of nuclear DNA synthesis was confirmed visually by autoradiography in [^3^H]-TdR labeled cultures treated with 2 x 10^−4^ M AAF (**Figure 3A**; labeling indices « 4%). In contrast, 10-fold higher labeling indices of « 40% were observed in the untreated control cultures (**Figure 3B**). No inhibitory effects of 1% v/v EtOH were observed, eliminating artifacts of the AAF-vehicle. None of these treatments caused morphological damage (Howard *et al.*, 1981).

**Figure 3.**
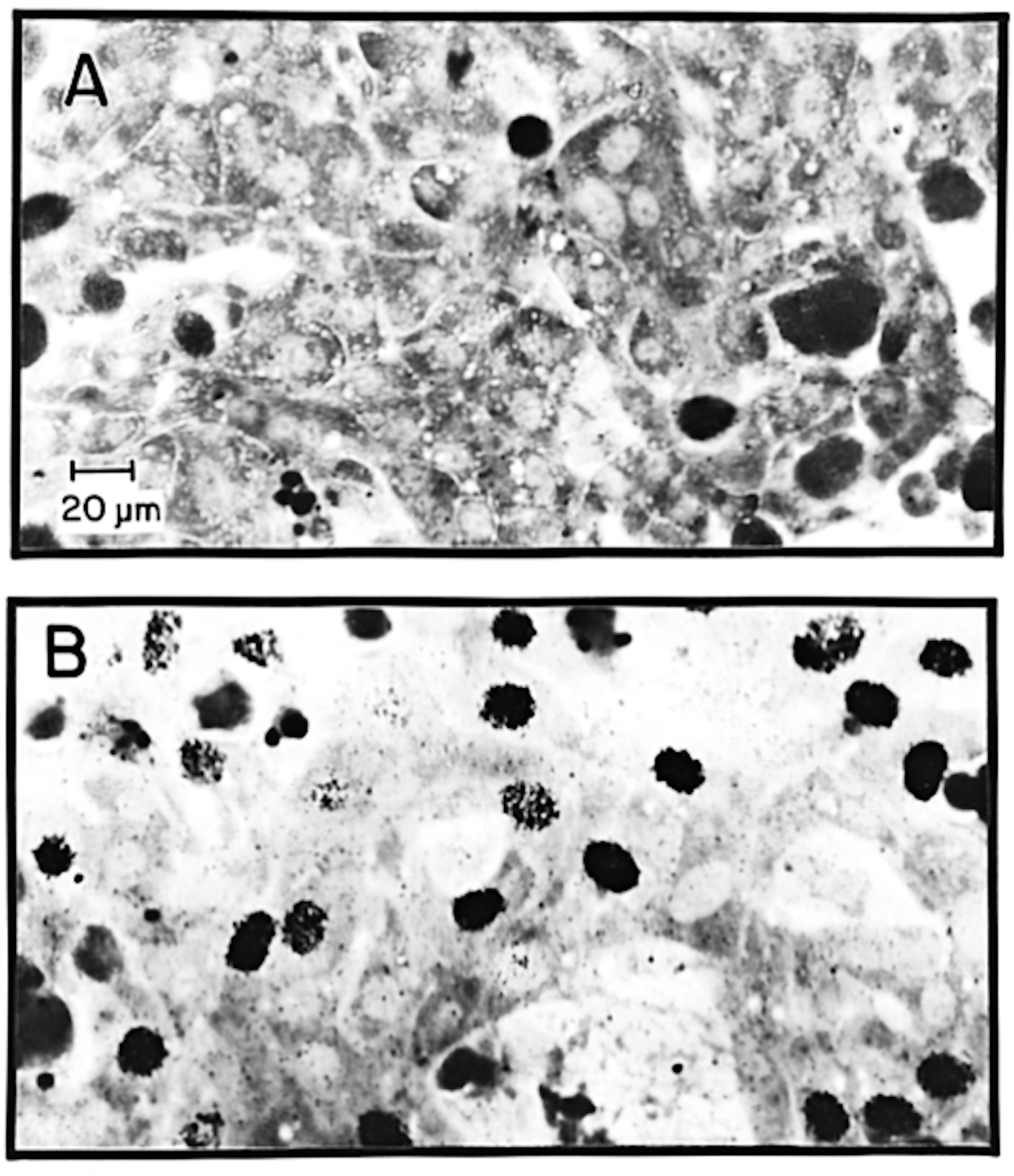
Autoradiography following growth initiation of quiescent hepatocytes. Cultured hepatocytes were plated from Fischer 344 rats. Unlabeled AAF (2 x 10^−4^ M, panel **(A)**) or 1% v/v EtOH (panel **(B)**) were added to 12 day-old cultures fluid-changed to fresh media plus 50 ng each of insulin, glucagon and EGF/mL. The cultures were labeled with 1.25 μCi [^3^H]-TdR/mL (3 μM TdR) between 12-24 h, and prepared as described in Materials and Methods. Representative photomicrographs are shown of both treatment conditions. Specifically labeled hepatocytes display dark black grains in round or ovoid nuclei (**inset bar** in (**A**) = 20 μm). Amorphously shaped dark objects are crystal violet staining artifacts. (Modified from Leffert et al., 1983, and used by permission of Springer Publishing.)

### Scatchard and growth cycle analyses of [^14^C]-AAF binding

When curves of the binding of [^14^C]-fluorene residues to cells in hepatocyte cultures were expressed as Scatchard plots, two classes of macromolecular binding targets – designated Site I and Site II – were observed in 4 day-old cultures (**Figure 4, main panel**). The source data are shown in the **inset** to **Figure 4**.

**Figure 4.**
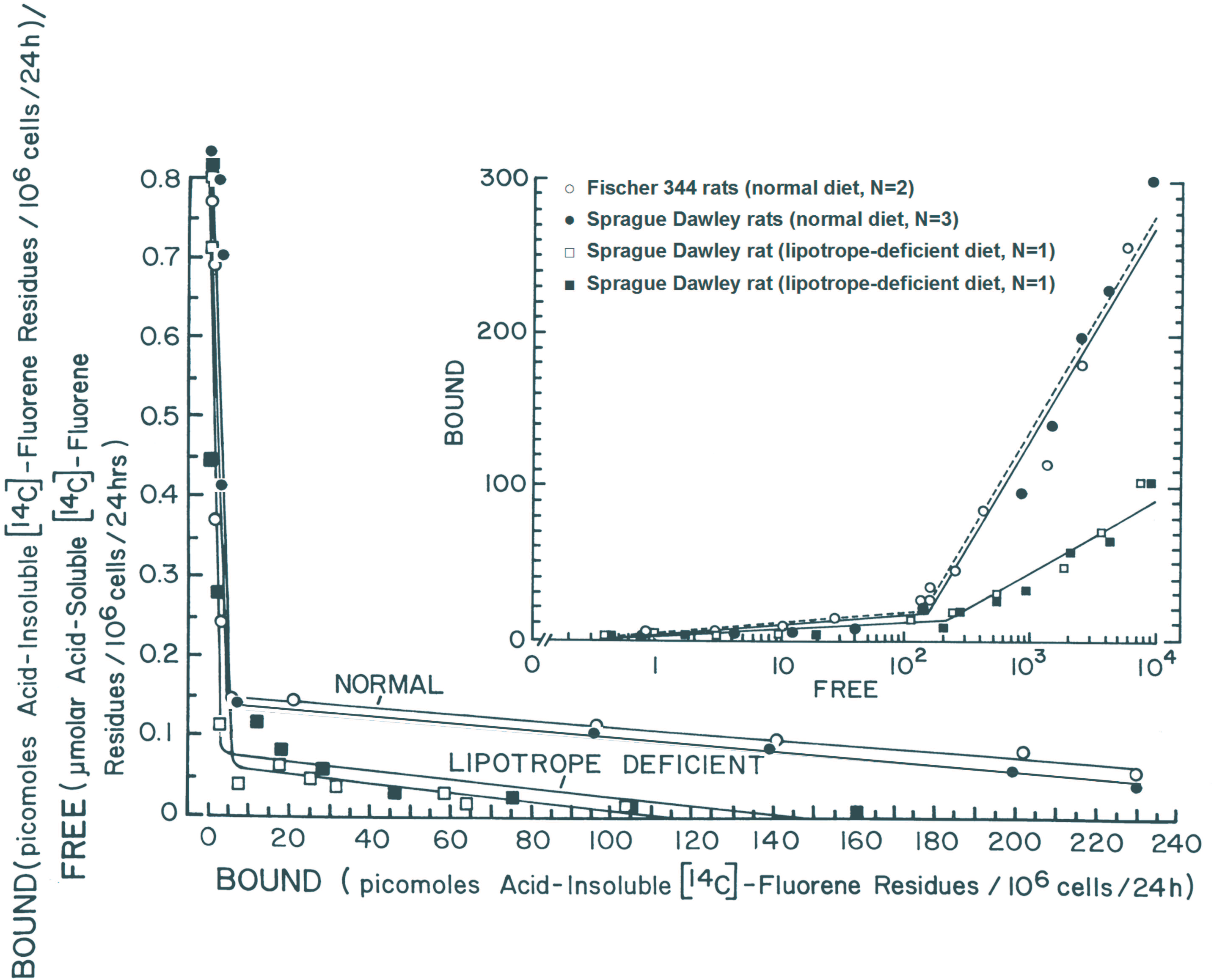
Quantitative binding of [^14^C]-AAF reveals two classes of AAF macromolecular binding sites. Four days post-plating, cultured adult rat hepatocytes from chow-fed Fischer 344 (**O** [each point is the average of plated cells from 2 individual rats]) or SD rats (• [each point is the average of plated cells from 3 individual rats]), or from 2 individual lipotrope-deficient SD rats plotted separately (□, ■), were isolated after 24 h labeling with [^14^C]-AAF over a wide concentration range: 1.25, 3.1, and 6.3 x 10^−7^ M; 1.25, 3.1, and 6.3 x 10^−6^ M; 1.8, 2.5, 3.9, and 6.7 x 10^−5^ M; and 1.2, 2.3, and 4.6 x 10^−4^ M. The **inset** figure shows experimental results: y-axis, BOUND (picomoles acid-insoluble [^14^C]-fluorene residues/10^6^ cells/24 h); x-axis, FREE (μmolar acid-soluble [^14^C]-fluorene residues/10^6^ cells/24 h). The results were re-plotted in Scatchard-like form (**main panel**): y-axis, BOUND [(picomoles Acid-insoluble [^14^C]-fluorene residues/10^6^ cells/24 h)/FREE (μmolar Acid-soluble [^14^C]-fluorene residues/10^6^ cells/24 h)]; x-axis, BOUND (picomoles Acid-insoluble [^14^C]-fluorene residues/10^6^ cells/24 h). Briefly, the K_D_ = −1/ [(slope of the curve)] = −1/([net Δ in B/F/net change in B]); and, the B_MAX_ = x-axis intercept. Further details including derivative constants are given in the text and elsewhere (Leffert *et al.*, 1983).

The values of the apparent affinity and apparent binding capacity constants obtained for Site I were similar among chow-fed Fischer 344, chow-fed Sprague Dawley (SD), and lipotrope-deficient SD hepatocytes: K_D[APPARENT]_ ≈ 5 x 10^−6^ M; and, B_MAX[APPARENT]_ ≈ 5.6 pmols/10^6^ cells/24 h. The Site II K_D[APPARENT]_ was also similar among all 3 groups (K_D[APPARENT]_ ≈ 1.5 x 10^−3^ M), but 1000-fold > the Site I affinity constant. On the other hand, although the Site II B_MAX[APPARENT]_ constant of cultured hepatocytes from both chow-fed strains, Fischer 344 and SD, was ≈ 350 pmols/10^6^ cells/24 h, the B_MAX[APPARENT]_ of the lipotrope-deficient SD hepatocytes was ≈ 126 pmols/10^6^ cells/24 h. These results suggested that the Site II B_MAX[APPARENT]_ constant might be sensitive to dietary conditions, which are known to affect hepatocyte growth properties (Leffert and Koch, 1977). In addition, the results of [^3^H]-AAF autoradiography of hepatocytes treated with 2 x 10^−5^ M AAF (10-fold higher than the Site I K_D[APPARENT]_) indicated that non-parenchymal cell contaminants were unlikely to have accounted for these binding differences (Koch *et al.*, submitted, 2017).

Plots of derivative constants determined by Scatchard analyses of acid-insoluble bound and acid-soluble free fluorene residues, following incubation over a 12-day time course with a wide concentration range of [^14^C]-AAF (1.25 x 10^−7^ M – 4.6 x 10^−4^ M), revealed a growth cycle dependence of Site I and Site II constants in cultured hepatocytes from chow-fed Fischer 344 rats. Between days 2-5 post-plating, concomitant decreases (≈ 30 fold) occurred in the Site I B_MAX[APPARENT]_ and Site I K_D[APPARENT]_ (**Supplementary Figure 1A and Supplementary Figure 1B**, respectively). Thereafter, both curves remained constant at their lower levels through day 12. In contrast, between days 2-5 post-plating, both Site II parameters increased 2-3 fold (B_MAX[APPARENT]_ and K_D[APPARENT]_, **Supplementary Figure 1A and Supplementary Figure 1B**, respectively). Thereafter, both curves remained constant at their higher levels through day 12. Site I and Site II values from the independent experiments were in good agreement; yet none of the curves followed the U-shaped patterns observed for constitutive growth cycle dependent differentiated functions in this system (Leffert *et al.*, 1978).

### [^14^C]-AAF and [^3^H]-AAF binding to genomic DNA

Three day-old log phase cultures were exposed to varying extracellular concentrations of [^14^C]-AAF (6 x 10^−8^ M – 2 x 10^−5^ M; **Figure 5A**). Twenty-four h later (day 4), the binding of acid-insoluble [^14^C]-fluorene residues to purified genomic DNA was measured as a function of acid-soluble [^14^C]-[AAF]_i_ (2 x 10^−7^ M – 4 x 10^−4^ M; Figure 5B). The curve of [^14^C]-fluorene binding to DNA showed saturation with a B_MAX[APPARENT]_≈ 6 pmols/10^6^ cells/24 h, and a K_D[APPARENT]_≈ 2 x 10^−6^ M, as determined in **Figure 5B**. The values of these constants were consistent with the Site 1 values of K_D[APPARENT]_ and B_MAX[APPARENT]_ determined by Scatchard analyses in **Supplementary Figure 1 and Figure 4**. Under similar culture conditions, starting with purified nuclei harvested from 4 day-old hepatocyte cultures, [^3^H]-labelled fluorene residues preferentially formed covalent DNA adducts with high molecular weight [^3^H]-TdR-labeled genomic DNA fragments (> 6-40 kb), as shown by flat-bed agarose gel electrophoresis, fluorography and scanning densitometry (Leffert *et al.*, 1983).

**Figure 5.**
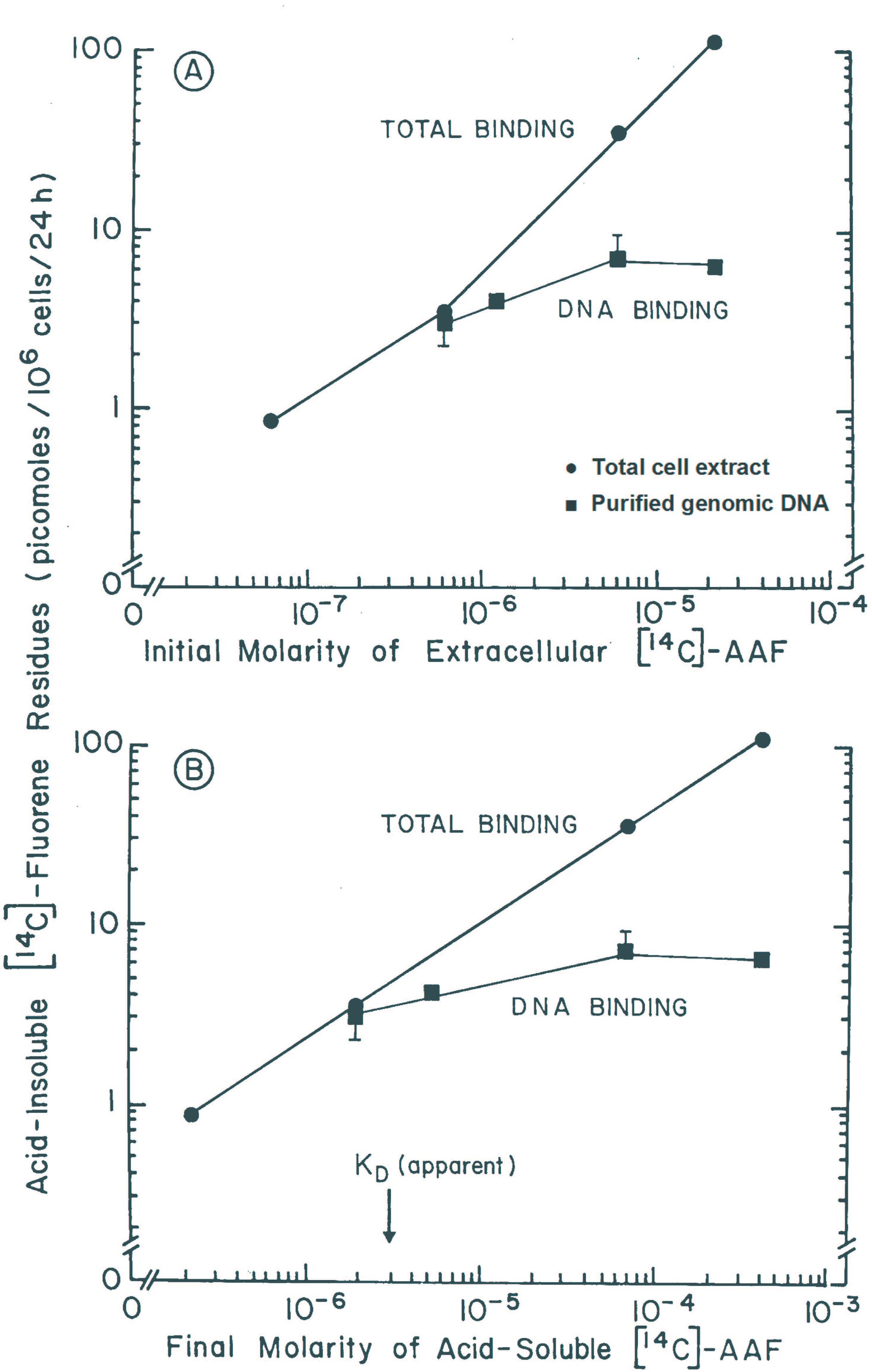
Saturable binding of [^14^C]-AAF to genomic DNA from cultured hepatocytes. Hepatocytes were plated from Fischer 344 rats. Purified genomic DNA (■) and total cell extracts (•) were isolated from 4 day-old cultures exposed on day 3 to a 4-log concentration range of [^14^C]-AAF, as described in Materials and Methods. **(A)** y-axis, Acid-insoluble [^14^C]-fluorene residues (picomoles/10^6^ cells/24 h), as a function of the initial molarity of [AAF]_o_ (6 x 10^−8^ – 2 x 10^−5^ M, x-axis). **(B)** y-axis, Acid-insoluble [^14^C]-fluorene residues (picomoles/10^6^ cells/24 h), as a function of the final concentration of acid-soluble [^14^C]-[AAF]_i_ (2 x 10^−7^ – 4 x 10^−4^ M, x-axis).

Parallel cultures were labeled for assays of total TCA-insoluble binding (**Figure 5**). In contrast to saturable genomic DNA binding, total [^14^C]-AAF binding to TCA-insoluble cellular material yielded linear curves, which did not saturate over the examined ranges of [^14^C]-AAF concentration (6 x 10^−8^ M – 4 x 10^−4^ M), as shown in **Figure 5A** and **Figure 5B**, respectively. These results suggested that hepatocytes contained additional pools of macromolecular binding targets, consistent with the Site II constants.

### [^3^H]-AAF binding to proteins

The B_MAX[APPARENT]_ for Site II_DAY 4_ (**Figure 4**) was ≈ 350 pmols/10^6^ cells/24 h or 2.6 x 10^8^ sites/cell, ≈ 100-fold higher than the B_MAX[APPARENT]_ associated with genomic DNA, the so-called Site I_DAY 4_ constant. This observation suggested that Site II was associated with a much larger pool of non-DNA binding targets. This was investigated by labeling 3 day-old cultures for 24 h with 2 x 10^−5^ M [^3^H]-AAF. Culture media and total cellular lysates were analyzed by SDS-PAGE and fluorography (Leffert *et al.*, 1983; Kruijer *et al.*, 1986). Densitometry scans of the fluorograms over the 24 h time course are shown in **Figure 6**. Binding of [^3^H]-fluorene residues to specific proteins in the media (**top tracing**: Mr ≈ 80, 68, 59, 27, and 13.7 kDa) and cytosols (**bottom tracing**: Mr ≈ 56 and 31 kDa) was observed; protein secretion was markedly delayed. The 68 kDa band, presumably albumin, was the most heavily labeled band in the media; however, the profile of the radiolabeled proteins in the media differed from that of the labeled cytosolic proteins (13-140 kDa; Leffert *et al.*, 1983). Pre-treatments of the media or the culture extracts with 50 μg pronase/mL (prior to SDS addition) led to the disappearance of all of the observed bands (data not shown). In separate cultures under similar conditions, phenobarbital pre-treatment also reduced the level of radioactivity incorporated into cellular proteins by ≈ 50%, but it did not alter the qualitative tracing pattern (data not shown); this reduction may have been due to changes in overall processing of AAF, such as increased conjugation of [^3^H]-AAF with glucuronic acid and reduced sulfate ester formation (Matsushima *et al.*, 1972). Neither media nor cytosolic labeling was detected by mixing fresh culture fluids or cellular lysates with [^3^H]-AAF, thereby eliminating artifacts of non-specific binding or extracellular protein synthesis. AAF binding specificity was suggested because different cytosolic labeling patterns occurred when cultures were incubated with the carcinogen [^14^C]-3-methylcholanthrene, for which most of the ^14^C was covalently bound to ligandin subunits, not to albumin (Leffert *et al.*, 1983).

**Figure 6.**
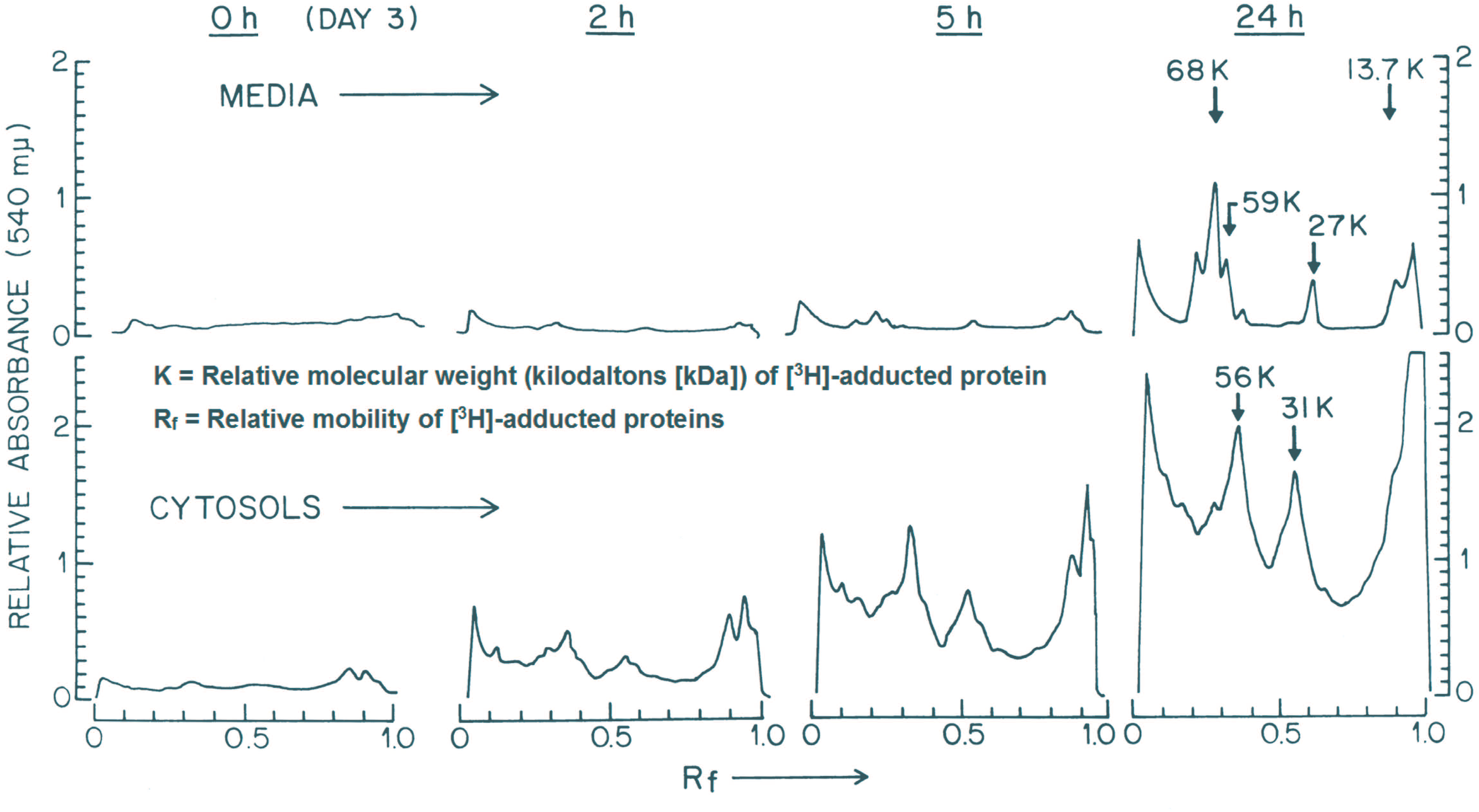
Densitometry scans of AAF-adducted cellular proteins from fluorograms of 10% SDS-PAGE gels. Hepatocytes were plated from Fischer 344 rats. Cultured cells were incubated with [^3^H]-AAF (2 x 10^−5^ M) at 3 days post-plating, and harvested for gel analysis as described elsewhere (Leffert *et al.*, 1983; Kruijer *et al.*, 1986) at the times indicated (h) along the top of the figure (K = kDa). Y-axes, Absorbance at 540 mμ; x-axes at each harvest time, R_f_ values. (**Top panel**) Proteins secreted into the media (albumin is prominently adducted; see ↓ at 68 kDa). (**Bottom panel**) Cytosolic proteins.

### [^14^C]-AAF binding to purified liver nuclei

Nuclei isolated from quiescent cultures contained NADPH-dependent enzymatic activities that, when incubated with 6.5 x 10^−6^ M [^14^C]-AAF (≈ Site I_DAY 4_ K_D[APPARENT]_), generated [^14^C]-fluorene residues bound to nuclear-associated DNA, RNA, protein and lipid targets (**Table 1**). Microsomal contamination was unlikely to account for the observed P450 activities since the outer nuclear membranes and endoplasmic reticulum were removed by Triton X-100 and shearing (Blobel and Potter, 1966). Non-specific binding (*i.e.*, when NADP replaced NADPH) was equivalent to no pre-treatment; it averaged 20.1 ± 4.4% (σ) of maximal NADPH-dependent binding. Thus, in the presence of NADPH, the percentages of maximal [^14^C]-fluorene residues bound to molecules in pre-treated samples were 100%, with washing, consistent with nuclear integrity; 23%, with boiling, suggestive of adduction to heat-denatured molecules; 57%, with DNase I treatment; 98%, with RNase I; and 35%, with pronase, suggestive of the presence of DNA, RNA and protein adducts, and of complexes inaccessible to any of these separate enzymes, respectively; and, 30%, following treatment with Triton X-100, suggestive of lipid adducts. The nature of the molecules that failed to survive individual pre-treatments was not determined.

**Table 1.**
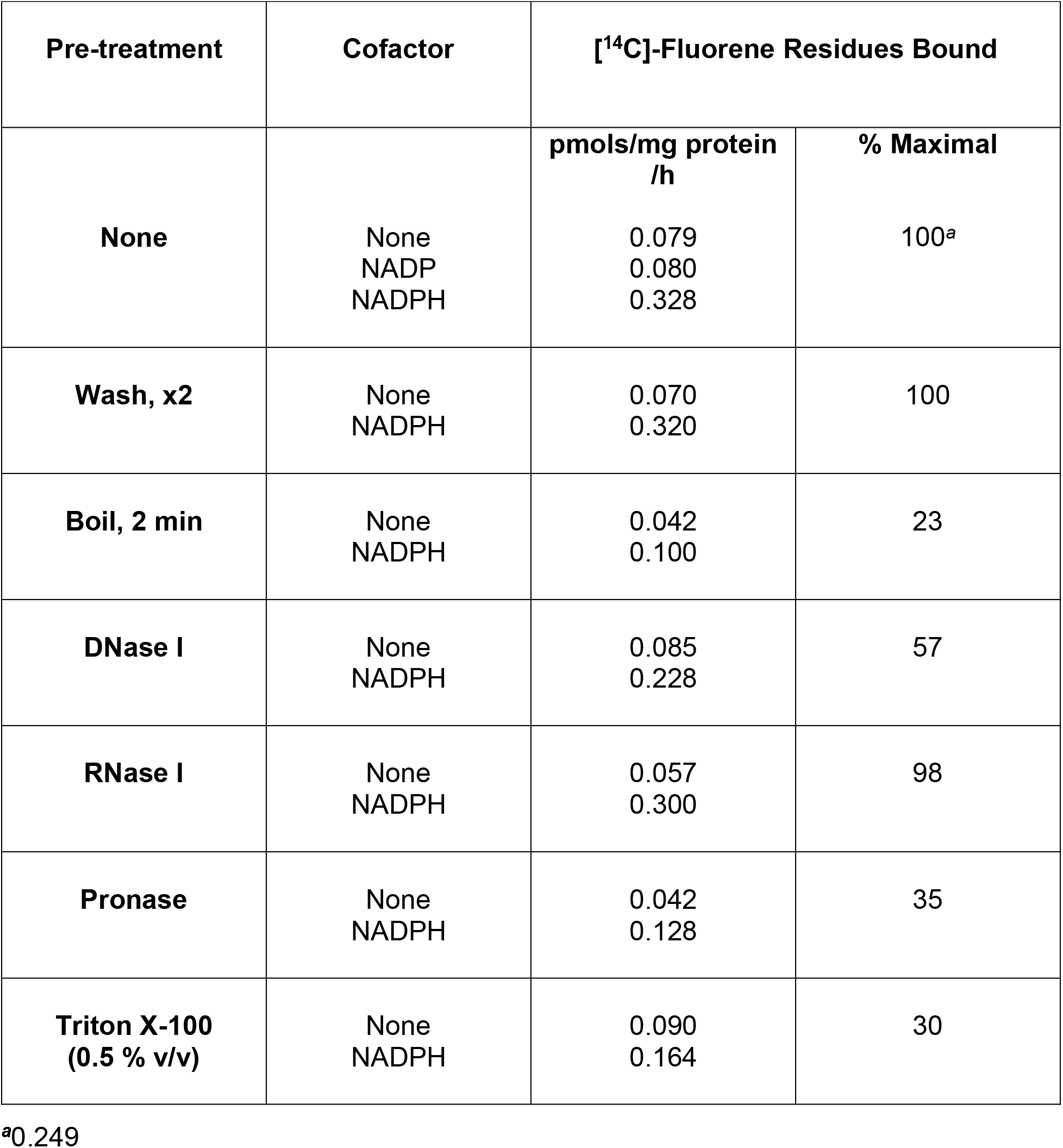
Covalent binding of [^14^C]-AAF to rat hepatocyte nuclei isolated from primary cultures of adult hepatocytes. Nuclei were isolated from washed 12 day-old quiescent Fischer 344 hepatocyte monolayer cultures, and subjected to various physicochemical or enzymatic pre-treatments (50 μg enzyme/mL). The recovered nuclei (1.4 mg DNA/mL) were incubated without or with 1 mg of a control (NADP) or cytochrome P450-specific cofactor (NADPH)/mL and [^14^C]-AAF (6.5 x 10^−6^ M) at 37°C for 1 h (Leffert *et al.*, 1983). The reactions were stopped by bringing the mixtures into 10% (v/v) TCA at 100°C for 5 mins. Insoluble material was precipitated onto Whatman GF/C™ filters, which were washed with ice cold EtOH, dried, counted to errors of ± 1%, and expressed as pmols/mg protein/h. The % **Maximal** value for each pre-treatment was calculated as follows: {([NADPH] – [None]) ÷ (0.328 – 0.079)} x 100. (Modified from Leffert *et al.*, 1983, and used by permission of Springer Publishing.)

## DISCUSSION

In this second of two reports (Koch *et al.*, submitted, 2017), investigations of AAF binding to cellular macromolecules were performed in long-term cultures of primary adult rat hepatocytes permissive of growth cycle and G_0,1_→S→G_2_→M transitions. Ploidy did not detectably affect AAF binding (Tulp *et al.*, 1978): proliferating hepatocytes were mononucleated (Leffert *et al.*, 1977; Koch and Leffert, 1979) and proliferation time courses lacked inflection points or multiple peaks suggestive of mixed ploidy.

Two classes of macromolecular AAF-binding sites were revealed by Scatchard analyses: a high-affinity low-capacity site; and a low-affinity high-capacity site. During the growth cycle, Site I B_MAX_ and K_D_ levels decreased, while Site II B_MAX_ and K_D_ levels increased. These alterations likely reflected growth-state and age-dependent changes in the differentiated status of nuclear and cytoplasmic P450 activities (specifically, N-2-hydroxylation), or changes in compartmentalized [AAF]_i_. Alternatively, since fluid changes were not made, media components regulating these constants may have become limiting *(e.g.*, hydrocortisone-succinate); or build-up of specific inhibitors may have occurred. The potential carcinogenic relevance of these inversely related growth state-dependent patterns of Site I and Site II expression remains to be determined.

[^14^C]-fluorene residues were bound covalently to RNA, DNA, protein and lipid in G_0_-hepatocyte nuclei, but these findings did not identify complexes associated with Site I or Site II. Several considerations suggested Site I is DNA-associated: [^14^C]-fluorene binding to genomic DNA required a Cyp1A2 protein, possessing a nuclear localization signal identified from primary sequence analysis (Koch *et al.*, submitted, 2017), associated with mitochondria-free nuclei (Blobel and Potter, 1966); [^14^C]-fluorene binding saturated DNA in log phase cells at 6 pmols/10^6^ cells/24 h, the Site I_DAY 4_ B_MAX[APPARENT]_, whereas, [^14^C]-fluorene binding in total cell extracts was linear over the same 2000-fold [AAF]_i_ range; at low [AAF]_o_ and [AAF]_i_ (6 x 10^−7^ M and 2-4 x 10^−6^ M, respectively), genomic DNA affinities for [^14^C]-fluorene residues were ≈ Site I_DAY 4_ K_D[APPARENT]_ determined by Scatchard analyses.

A binding rate of 3.6 million adducts/rat genome can be calculated from results based upon the Site I_DAY 4_ B_MAX_ (6 pmols [^14^C]-fluorene residues/10^6^ cells/24 h). This calculation assumes that: i) All hepatocytes bind AAF; ii) Diploid hepatocytes contain 8 pg nuclear DNA/cell (Bibbiani *et al.*, 1969; Blobel and Potter, 1966); iii) Diploid hepatocyte genomes contain 2.51 x 10^9^ bp ≈ 5 x 10^9^ nts (Twigger *et al.*, 2008); and, iv) [^14^C]-fluorene residues bind randomly 1:1 to accessible dG sites. Thus, Site I_DAY 4_ saturation led to 7.2 bound [^14^C]-fluorene residues/10,000 nts, in good agreement with the results of feeding 0.02% AAF to adult rats (Poirier *et al.*, 1990). Importantly, this calculation is restricted by the above assumptions; and, by current lack of knowledge of the influence, magnitude and effects of non-genic or genic, non-coding, and repetitive DNA on AAF adduction in living hepatocytes.

Notably, >50% of the human genome consists of repetitive tracts; many are G·C rich and essential to genome function (Shapiro and von Sternberg, 2005). Numerous repeats have non-B triplex and quadruplex structure (Wells *et al.*, 1988) which might hinder AAF-metabolite binding to dG residues (Veaute and Fuchs, 1991; Lindsley and Fuchs, 1994; Culp *et al.*, 1993; Savreux-Lenglet *et al.*, 2015) or misdirect it (*e.g*., N-acetoxy-2-acetylaminofluorene binding to unpaired adenine and cytosine residues in non-B DNA structures [Kohwi-Shigamatsu *et al.*, 1988]). AAF also induces B→Z-DNA transitions (Santella *et al.*, 1981); and, except at B-Z junctions, Z-DNA is more resistant to DNA nucleases than B-DNA (Rich *et al.*, 1984). Given this complexity, structural determinations of AAF-DNA adducts along these tracts will be needed for precise quantification of adduct levels/genome.

Unexpectedly, when quiescent hepatocyte cultures were stimulated to grow, [AAF]_o_ far above the Site I K_D[APPARENT]_ were required to inhibit DNA synthesis initiation, S-phase entry and ongoing DNA replication; none of these processes was blocked by 0.6 μM [AAF]_o_ (≈ Site I_DAY 12_ K_D[APPARENT]_). These findings indicated that saturation of genomic DNA with covalent adducts (Site I_DAY 12_ B_MAX[APPARENT]_ ~ 2 pmols [^14^C]-fluorene residues /10^6^cells/24 h) was insufficient to block G_0,1_→S→G_2_→M transitions.

How DNA synthesis proceeds in AAF adduct-saturated genomes in living hepatocytes is unknown. In cell-free systems, AAF adducts differentially affect DNA synthesis. For instance, primer extension studies using oligonucleotides containing AF- or AAF-adducted *Nar*I restriction sites, showed that, dependent upon DNA sequence context, phage and bacterial DNA polymerases stalled or bypassed single stranded (ss) sites containing dG-C8-AAF or dG-C8-AF adducts, respectively (Belguise-Valladier and Fuchs, 1995).

DNA synthesis initiation and chain elongation were also examined in eukaryotic extracts using DNA templates modified site-specifically with dG-C8-AAF adducts (Hoffman *et al.*, 1996). Here, the evidence suggested that replicative polymerase ε and α-primase (but not nucleotide excision repair [NER] polymerase β) bypassed adducted initiation sites in double-stranded forked templates (allowing initiation and continuous 5’→3’ elongation of leading strand synthesis), whereas pol δ activity stalled at adducted sites of chain elongation in ss-templates (blocking discontinuous 3’←5’ lagging strand synthesis of Okazaki fragments).

It is possible, therefore, considering both cell-free system and cultured hepatocyte findings together, that translesion synthesis (TLS) occurs in adduct-saturated genomes of proliferating hepatocytes exposed to submicromolar [AAF]_o_. Eukaryotic TLS employs less stringent polymerases (such as pol η, ι, ν and ζ) that substitute for more stringent leading strand polymerases operative during S-phase (Hedglin and Benkovic, 2017). Further investigations with primary hepatocyte systems, competent of forming DNA adducts during G_0,1_→S→G_2_→M transitions, will be necessary to identify structures and genomic distributions of adducts that bind to DNA at submicromolar [AAF]_o_; and, to identify DNA polymerases and mechanisms of DNA replication along adduct-saturated DNA.

While mechanisms underlying saturable binding of [^14^C]-fluorene residues to genomic DNA may still involve elevated NER, this explanation is hard to reconcile with findings that >10-fold higher [AAF]_o_ were required to induce DNA repair in other primary hepatocyte systems (Williams, 1977).^2^ Recent investigations reveal extensive complexity of global and transcription-coupled NER (Marteijn *et al.*, 2014); combined with cell-free DNA synthesis reports, and structural and sequence-context variables that dictate bypass and repair of dG-C8-AF, dG-C8-AAF and dG-N^2^-AAF adducts (Veaute and Fuchs, 1991; Lindsley and Fuchs, 1994; Culp *et al.*, 1993; Mu *et al.*, 2012), the saturable Site I binding properties of cultured long-term hepatocytes will almost certainly be regulated by many biophysical and biochemical factors.

Several observations suggested Site II is associated with proteins. AAF metabolites formed protein adducts, via nuclear and cytoplasmic P450s, with higher K_D[APPARENT]_ (1.5 x 10^−3^ M) and B_MAX[APPARENT]_ values (350 pmols/10^6^ cells/24 h) than Site I genomic DNA constants; [^3^H]-fluorene-adducted proteins were undetectable at [AAF]_o_ < 10^−6^ M (Leffert *et al.*, 1983). This might explain why [AAF]_o_ ≥ 2 x 10^−4^ M, nearing the Site II K_D[APPARENT]_, were required to block DNA synthesis initiation and replication, and markedly depress S-phase entry rates. These results suggested that [AAF]_o_, near the Site II K_D_, exerted effects indirectly by inactivating proteins or other molecules required for DNA, RNA and protein synthesis, and continued hepatocyte proliferation; and, by blocking intracellular protein trafficking and secretion (Harson and Williams, 1979). These conclusions are consistent with the delayed secretion of [^3^H]-fluorene-adducted proteins observed in log phase cultures; with the inhibition of DNA replication and prolonged S-phase entry observed in N-OH-AAF-injected rats 20 h after 70% partial hepatectomy (‘PH’), when 1^st^ rounds of DNA synthesis start (Koch and Leffert, 1979; Zurlo *et al.*, 1986); and, with prolonged G_0,1_→S phase transitions following aberrant cell cycle signaling, exemplified by elevated *p53* and *p21* expression in adult rats fed AAF for 7 days before 70% PH (Ohlson *et al.*, 1998). Elevated *p53* and *p21* oncogene expression are involved in DNA damage responses (DDR) that promote cellular senescence associated with irreversible G1- and G2-arrest (Gire and Dulic, 2015), and immune surveillance suppression of mouse hepatocellular carcinoma (HCC; Xue *et al.*,2007). Further experiments with long-term cultures can help identify cell cycle transition signaling proteins and time intervals inactivated or blocked by AAF, including possible hepatocellular senescence responses triggered only at [AAF]_o_ above the postulated TLS-associated Site I K_D_ or approaching the postulated DDR-associated Site II K_D_.

Several corollaries are suggested from observations made with this model system: i) If DNA adducts initiate HCC, then carcinogenic thresholds are much lower than concentrations inducing cultured rat and human hepatocyte DNA repair (Williams, 1977; Spilman and Byard, 1981; Strom *et al.*, 1983; Monteith *et al.*, 1988) or elevated human fibroblast mutation rates (Strom *et al.*, 1983); ii) Mutated hepatocytes might arise, absent lethal or growth selective environments, because genomes saturated with DNA adducts do not restrict growth transitions in living hepatocytes; and, iii) Site I B_MAX_ constants are quantitatively equivalent in hepatocytes from rats fed lipotrope-deficient or normal chow diets; by contrast, Site II B_MAX_ constants fall significantly in lipotrope-deficient compared to normal hepatocytes, suggesting a role for Site II in the promotion of HCC. This view is consistent with observations, prescient of human NAFLD-associated HCC (Haas *et al.*, 2016), that rats fed lipotrope-deficient diets manifest much higher than normal risks for HCC induced by AAF (Rogers, 1975); and, cultured lipotrope-deficient, lipid droplet laden hepatocytes behave like transformed cells (Leffert and Koch, 1977).

A critical question remains: As hepatocytes proliferate *in vitro*, can one or more type of precursor cell (Koch and Leffert, 2015), associated with altered Site I and/or Site II expression, emerge from AAF treatment to give rise to HCC? Continued investigations with this AAF processing and adduct-forming system may provide an answer to this most significant question, particularly if the longevity of this system^3^ can be extended to time intervals required for AAF to induce HCC *in vivo* (Teebor and Becker, 1971).

## ACKNOWLEDGMENTS

We thank Hal Skelly for additional technical assistance.

## FOOTNOTES

1 The primary culture systems we developed in the 1970s’ enable primary hepatocyte survival, differentiation (*e.g.*, the definitive marker of arginine biosynthesis, ornithine transcarbamylase) and proliferation from any inbred or outbred vertebrate source. In this manuscript, sustained and concomitant adult hepatocyte differentiation (AAF metabolism and macromolecular adduct formation) and proliferative capacity (parasynchronous growth cycles demonstrating lag, log and stationary phase through day 10, as well as synchronous G_0,1_→S→G_2_→M transitions in 12 day-old stationary phase cultures) were required to perform the investigations described here. These conditions of hepatocellular proliferation were not reported in short-term rat^2^ or longterm human^2^ hepatocyte cultures (Ullrich *et al.*, 2007).

2 Saturable AAF binding to DNA was not reported in short-term rat and human hepatocyte cultures (Spilman and Byard, 1981 [2-5 days]; Strom *et al.*,1983 [4-24 h]; Monteith *et al.*, 1988 [1-4 days]), yet quantitative [^14^C]-fluorene binding levels described in this paper overlapped with results from these other systems (for instance, 0.16 nmols fluorene/mg DNA/24 h at 6 μM [AAF]_o_ is a concentration close to the here-determined Site I_DAY 4_ K_D[APPARENT]_). To resolve the contrasting observations regarding saturation, uniform hepatocyte culture conditions and identical kinds of measurements are needed for comparison.

3 Primary hepatocytes described here have survived >48 days *in vitro* (not shown).

---

**Supplementary Figure 1. Changes in the Site I and Site II B_MAX[APPARENT]_ and K_D[APPARENT]_ constants during the growth cycle**. Hepatocytes were plated from Fischer 344 rats. The binding and uptake of [^14^C]-AAF were measured after 24 h of daily pulse-labeling with concentrations of 6 x 10^−8^ M, 6 x 10^−7^ M, 6 x 10^−6^ M, and 2 x 10^−5^ M [AAF]_o_, over a 12-day period post-plating. Scatchard analyses were performed and derivative constants determined by standard procedures as described in Materials and Methods, and in Figure 4. Site I (•) and Site II (■) constants in panels **(A) B_MAX[APPARENT]_** (Binding Capacity; y-axis), and **(B) K_D[APPARENT]_** (y-axis), were plotted as a function of time (x-axis).

